# DisDock: A Deep Learning Method for Metal Ion-Protein Redocking

**DOI:** 10.1101/2023.12.07.570531

**Authors:** Menghan Lin, Keqiao Li, Yuan Zhang, Feng Pan, Wei Wu, Jinfeng Zhang

## Abstract

The structures of metalloproteins are essential for comprehending their functions and interactions. The breakthrough of AlphaFold has made it possible to predict protein structures with experimental accuracy. However, the type of metal ion that a metalloprotein binds and the binding structure are still not readily available, even with the predicted protein structure. In this study, we present DisDock, a physics-driven deep learning method for predicting protein-metal docking. DisDock takes distogram of randomly initialized protein-ligand configuration as input and outputs the distogram of the predicted binding complex. It combines the U-net architecture with self-attention modules to enhance model performance. Taking inspiration from the physical principle that atoms in closer proximity display a stronger mutual attraction, this predictor capitalizes on geometric information to uncover latent characteristics indicative of atom interactions. To train our model, we employ a high-quality metalloprotein dataset sourced from the Mother of All Databases (MOAD). Experimental results demonstrate that our approach outperforms other existing methods in prediction accuracy for various types of metal ions.

## Introduction

Comprehending the interactions between metals and proteins is crucial for grasping the intricate molecular mechanisms underlying physiological processes. Proteins that bind to metal ions, known as metalloproteins, perform many functions within cells, including the storage and transport of metal ions (1). Moreover, it is noteworthy that approximately 38% of entries in the Protein Data Bank (PDB) involve at least one metal ion, and nearly half of all enzymes depend on specific metal associations for their functionality (2; 3). Since the functions of proteins are determined by their structures, understanding the structures of protein-metal complexes will provide scientists with a deeper insight into the functional roles and mechanisms of action of metalloproteins. Experimentally solving protein structures is highly time- and resource-consuming and can pose significant challenges for certain proteins. Consequently, researchers have turned to computational approaches to determine protein structures, achieving significant successes in recent years (4). Previous studies on the computational prediction of metalloprotein structures can be classified into two types.

The first type of study involves sequence- and structure-based methods that primarily focus on predicting the protein residues that bind to metal ions (5; 6; 7). These approaches utilize prior knowledge to design amino acid features. Sequence-based methods incorporate composition, position, structural, and physiochemical information as features. On the other hand, structure-based methods utilize features derived from structural motif analogies, such as backbone geometry and 3D arrangement of amino acids. These pre-designed features then served as input for machine learning or deep learning models to identify binding residues in proteins. While these methods have achieved high accuracy in predicting potential binding residues, few of them can reconstruct the three-dimensional structure of metal-protein complexes.

The second kind of method is designed to predict metal ion binding location while providing the metal-protein complex structure (8; 9; 10). One of the current state-of-the-art predictors for metal location is BioMetAll (11), which creates a spherical grid of equidistributed metal probes that embed the entire protein. Then, probes that satisfy several constraints are kept and grouped based on coordinating amino acids. However, it cannot guarantee the existence of a solution when a search region actually contains a metal ion. Another method MIB2 (10) constructs metal ion-binding templates, and utilizes a fragment transformation method to assess the structural similarity between these templates and a given protein structure. The alignment score ranks each prediction, and only those exceeding the set threshold are retained. Despite its advancements, MIB2 still has certain limitations, including reliance on input data quality, considerable computational time for processing, and the inability to carry out a batch of local searches using the web server.

Two newly-developed methods, Metal1D and Metal3D (12) were shown to improve the location prediction of zinc ions in protein structures using coordination motifs and 3D convolutional neural networks, respectively. However, structures larger than 3000 residues were not considered, and the models were trained solely on zinc ions, which limits their practical applications.

In this paper, we introduce DisDock, a distance-based model that incorporates the U-Net architecture (13) with self-attention modules to predict protein-metal docking. Inspired by the physical principle that atoms in closer proximity exhibit stronger mutual attraction, this predictor leverages input distances to extract hidden features representing the physical interactions among atoms. Importantly, by design DisDock has the potential to accommodate the flexibility of both ligands and proteins. In this study, we focus on docking metal ions to rigid protein structures for 16 metal ions commonly seen in protein structures.

The rest of this paper is structured in the following way: Section 2 provides details on our dataset and preprocessing procedure. Section 3 outlines the development of our training pipelines, model architectures, and postprocessing procedure. Section 4 presents the main results, focusing on docking accuracy. Finally, in Section 5, we summarize the paper and discuss potential directions for future work.

## Data

In this study, we focus on a dataset called the Mother of All Databases (MOAD) (14; 15; 16), a subset of the Protein Data Bank (PDB) (17). This dataset has the largest ligand-protein binding collections that involve protein crystal structures with a resolution of at least 2.5 Å to clearly identify biologically relevant ligands. In total, there are 331,372 protein-metal binding structures.

### Data Selection

To obtain valid results, our study restricts to high-quality protein-ligand pairs by excluding invalid raw data, such as proteins with missing parts or unknown amino acids. That is, we only focus on metal ions that are investigated in AlphaFill (18), MIB2 (10), or MetalPDB (19). As shown in Table S2 Table, we included 16 types of metal ions in our study: *Mg*^2+^, *Zn*^2+^, *Ca*^2+^, *Na*^+^, *Mn*^2+^, *K*^+^, *Fe*^3+^,*Ni*^2+^, *Co*^2+^, *Fe*^2+^, *Cu*^2+^, *Ba*^2+^, *Mo, W* ^6+^, *Cu*^1+^, and *Cd*^2+^. To prevent information leakage between training and testing datasets, similar pairs are grouped as closely as possible to stay within either the training or test part. we start by using T-Coffee (20; 21) to align protein sequences with structural information and calculate similarity scores between pairs. We then employ the Farthest point algorithm for hierarchical clustering to update distances between clusters and combine sequences that are alike. Our objective is to ensure that the initial observations in each cluster have a dissimilarity value of 0.3 or lower. Finally, we randomly merge clusters until we have three parts of the data, namely, training, validation and test, with a proportion of 70%, 10%, and 20% of the total pairs, respectively. S2 Table in the Supporting information provides an overview of the occurrence rate for each metal ion.

### Preprocessing

#### Pairwise distance matrix

In this study, we assume protein structures are rigid bodies. Given 3D coordinates for ligand of interest, we define a local region containing the closest 255 heavy atoms in the protein structures and compute a symmetric pairwise distance matrix D with size 256 × 256 for a metal ion and the chosen 255 protein atoms in the local region. In Figure 2, we show the Euclidean distance between the real metal binding location and the farthest atom among its closest ones in protein that ranges from 10 Å to 50.7 Å with median being 11.9 Å. The environment with 255 atoms strikes a good balance between including enough information about protein-metal binding and reasonable computational complexity. In mFASD (22), atoms within 5 Å of the bound metal are considered to be its neighborhood, and Wang’s method (23) instructs a probe to move within a box of size 18 Å × 18 Å × 18 Å.

An example of an input data point is shown in the left panel of Figure 1, where the binding location of a magnesium ion is to be predicted. The left plot represents the distance matrix with all diagonal elements being 0. Two green bars indicate distances between *Mg*^2+^ and local atoms in the protein. The blue region shows pairwise distances among atoms on the corresponding protein structure. Because each metal ligand only has one atom, the size of *D* is 256 × 256. For the current study where proteins are fixed as rigid objects, only the distances between the metal and protein atoms (the green region) need to change during the training process and distances among the protein atoms (blue region) are invariant.

**Fig 1.**
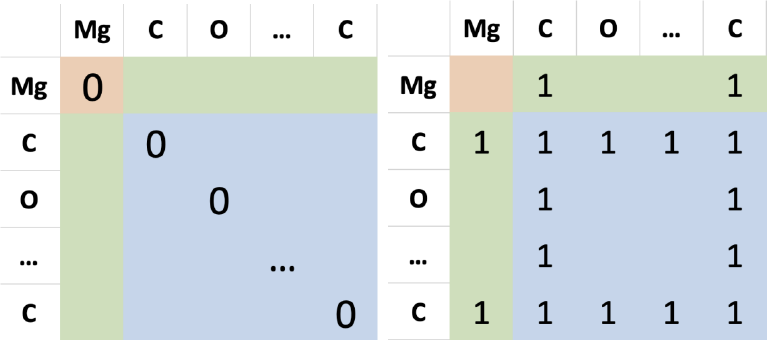
Left: The pairwise distance matrix D. Right: Indicator channel for Carbon. There are 21 channels that signify the atom types of proteins, determined by the specific residue and atom types they correspond to. Additionally, there is a separate channel dedicated to representing ions. Thus, they form inputs of size 23 × 256 × 256.

**Fig 2.**
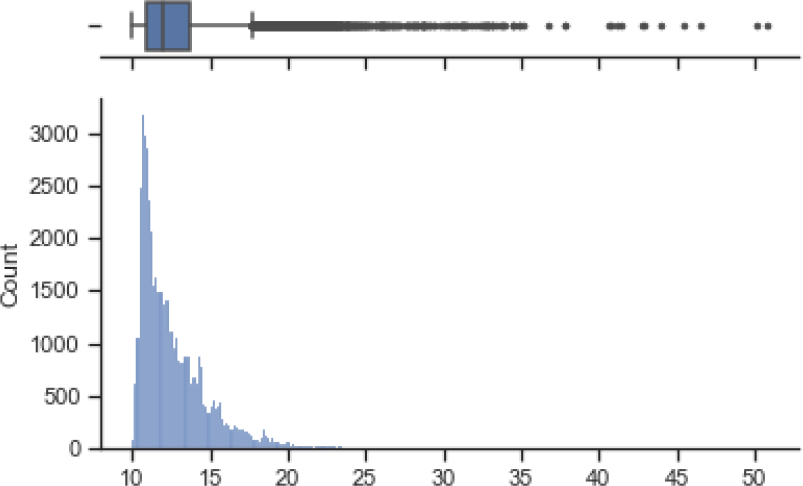
The distribution of Euclidean distances between metal ion to the farthest atom within a 255-atom local region. The majority of distances are less than 18 Å.

An example of atom type channels, a channel for the C atom of amino acids, is shown in the right panel of Figure 1. A value of 1 is assigned to any row or column corresponding to the C atom, and a value of 0 to all other positions. The encoding of each atom is based on its amino acid residue and type, and additional information can be found in Supporting information S1 Table. The input consists of a distance matrix on the first channel, followed by 21 channels indicating atom types, and one channel representing the metal ion. The resulting output has the size of 23 × 256 × 256. To generate inputs, we simulated initial metal ion locations using Algorithm 1 to mimic 3D random walk affected by a large temperature parameter *T* to ensure a random initial state. Then, given rigid proteins and simulated metal ion initial locations, we produce corresponding input boxes of size 23 × 256 × 256 for each ligand-protein pair.

#### Data Augmentation

Our goal is to predict the protein-metal complex of a protein and its metal ligand by moving the metal ion to its binding position in a chosen local region of the protein (a subset of atoms close to a location on the protein surface). In practice, the local region is not known since the binding position of the metal is not known. A plausible approach to address this is to predict the local region (or binding residues) first, and then predict the binding position of the metal with the predicted local region. As the predicted local region may have various degrees of overlap with the true local region, to simulate this situation of non-ideal local regions, we select multiple local regions from each protein-metal complex by varying the degree of overlap of selected local regions with the true local region. We expand our testing data by first randomly selecting five atoms from the 30 closest protein atoms to the metal ion. Using each of these neighboring atoms as a center, we then select 255 atoms closest to that center to form the local region.

##### Algorithm 1 Generating initial location

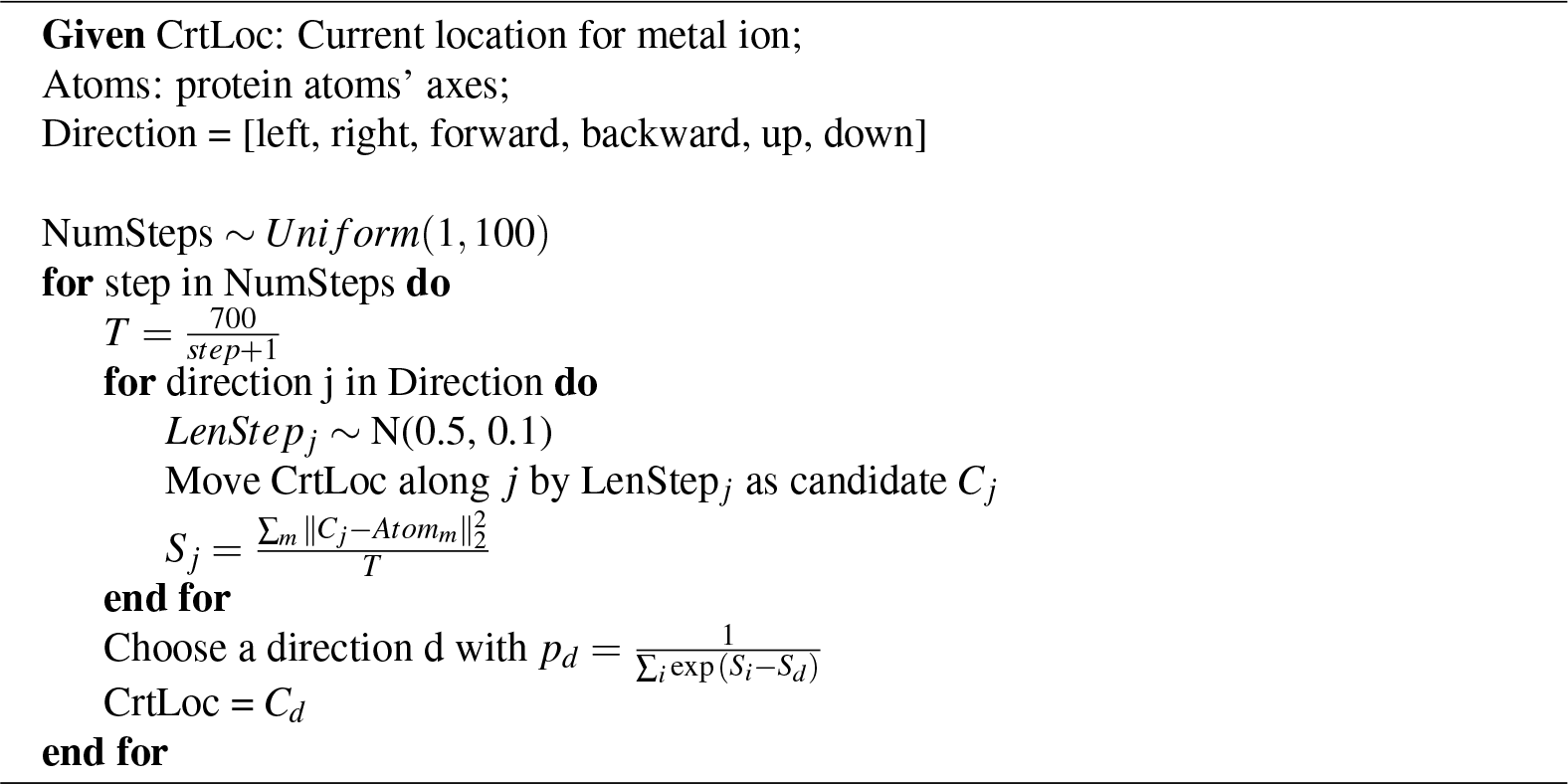

## Methods

The process of predicting protein-metal ion binding locations is presented in Figure 3 (a). The metal ion-protein complex is represented as a matrix of size 23 × 256 × 256. Then the DisDock model predicts a pairwise distance matrix with size 1 × 256 × 256, and the estimated metal ion coordinates are provided after the postprocessing procedure.

**Fig 3.**
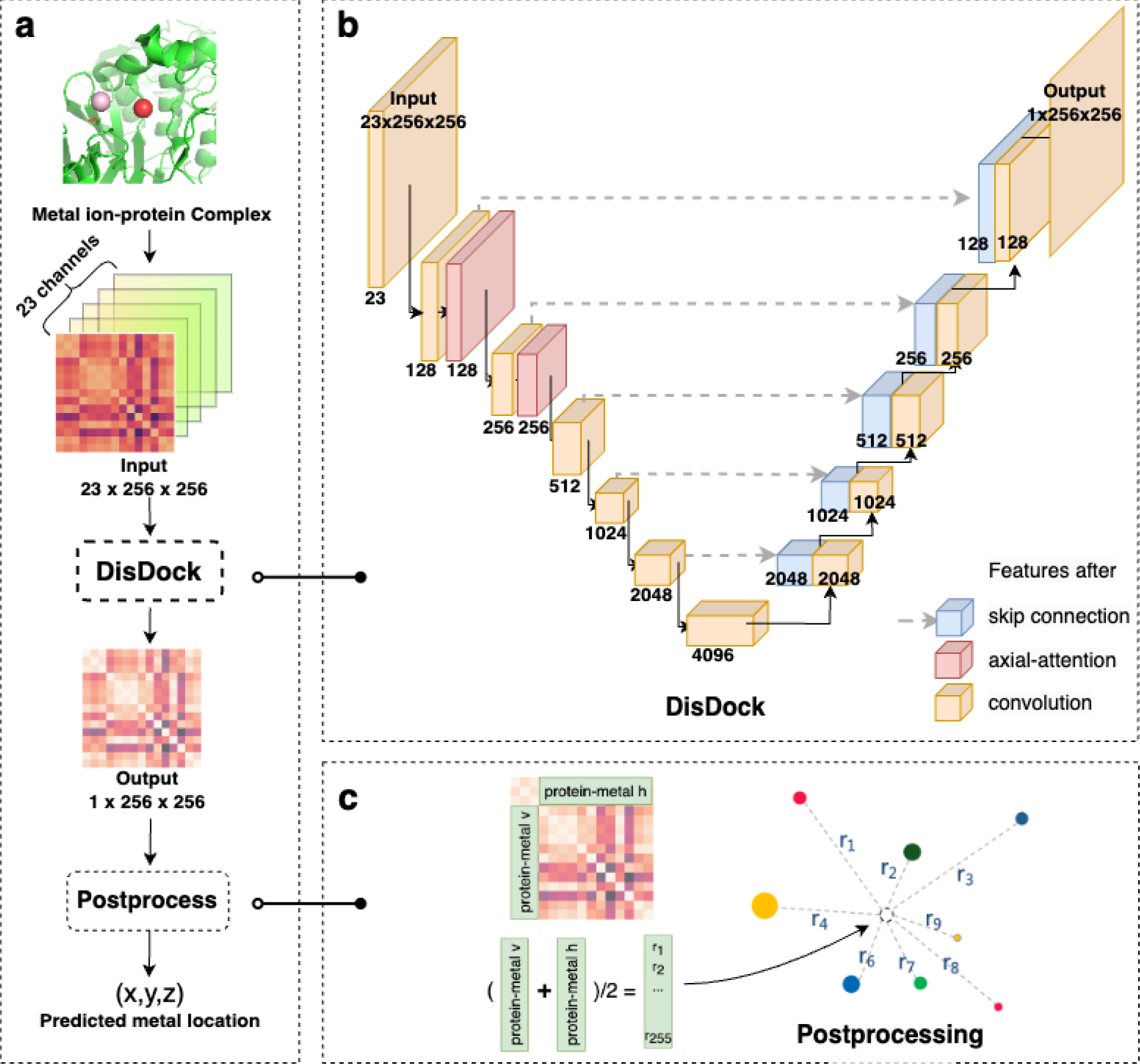
The overview of the redocking procedure. In the metal ion-protein complex, the true location of the metal ion is marked in red, while the randomly generated metal ion is represented in pink within the green protein environment. The complex is then encoded as an input including the relative distance between the actual protein and the initialized metal ion (pink), as well as the types of atoms in the local pocket environment. The DisDock model is used to learn atom-atom protein-metal ion interaction. Eventually, the predicted interaction is processed to estimate the metal ion location.

### Network Architecture

Prior work commonly employs an encoder-decoder network (24), comprising downsampling layers followed by a bottleneck layer and subsequent upsampling. Given the significant overlap of input and output distance matrices, direct information transfer offers notable benefits. In light of the premise, our foundational model is inspired by the ‘U-net’ architecture (25), implementing skip connections that interlink the *i*-th and (*n* − *i*)-th layers (with *n* layers total) to augment information dissemination. Alongside these connections, the ‘U-net’ structure integrates two modules: *C*_*k*_ for gradual downsampling and *CD*_*k*_ for upsampling. The former employs Convolution-Instance Norm-Leaky ReLU layers with k filters, while the latter employs Convolution-Instance Norm-Dropout-ReLU layers, also with *k* filters. Additional details can be found in the GitHub repository (26).

#### Axial-attention module (28)

Ever since its inception, the attention mechanism has gained widespread utilization for capturing and encoding long-range interactions, leading to numerous instances of cutting-edge performance (29; 30; 31). Nonetheless, the utilization of global self-attention encounters limitations when dealing with high-resolution images due to its computationally intensive nature, as it necessitates calculating relationships between every pixel and all others. A solution to this predicament involves decomposing 2D self-attention into two separate 1D self-attentions. This method is exemplified through the integration of axial attention module denoted as *A*_*k*_, with *k* denoting the number of filters. The architecture of this self-attention module draws inspiration from (27). As showcased by Figure 4, each axial-attention block encompasses a convolutional layer, succeeded by two consecutive axial-attention blocks applied sequentially along the height and width axes. Subsequently, the outputs undergo an additional convolutional layer, followed by the integration of a skip connection to amplify performance. Further details are accessible at GitHub repository (32).

**Fig 4.**
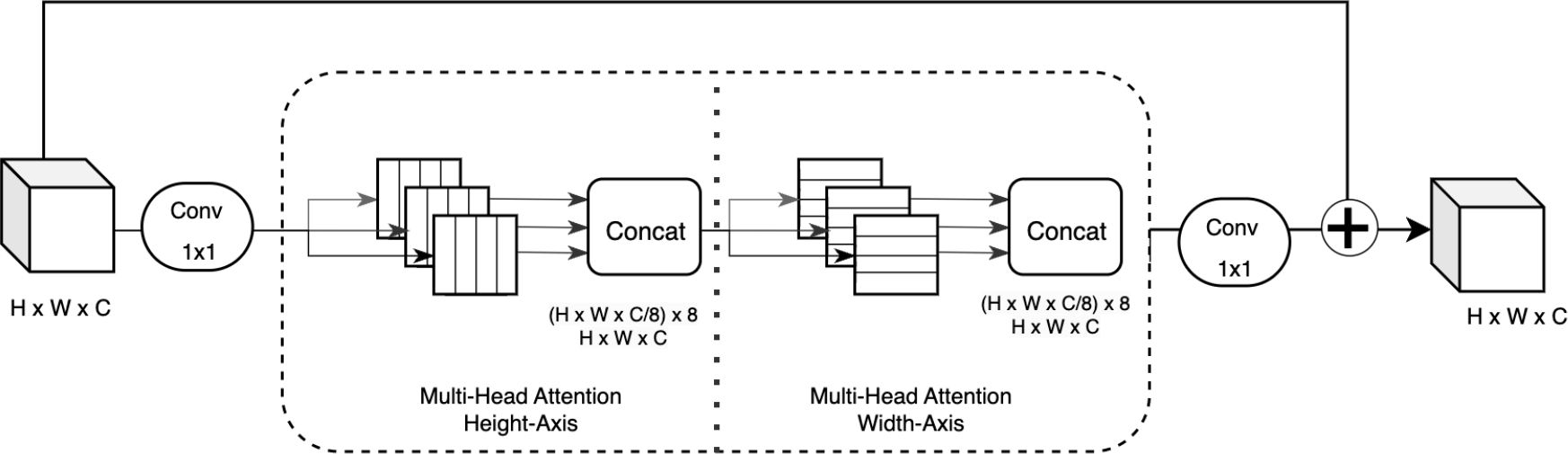
The axial-attention block comprises two successive axial-attention layers, which are applied sequentially along the height and width axes. The input is directed into these axial-attention layers, and the ensuing output undergoes processing by a convolutional layer to reorganize the latent features. To facilitate seamless information and gradient propagation during training, a skip connection is employed. Notably, the model integrates 8 attention heads. The illustration is derived from the architectural diagram presented in the paper “Axial-DeepLab: Stand-Alone Axial-Attention for Panoptic Segmentation” (27).

#### Atom-atom U-net with attention (DisDock)

In order to take more information from protein atoms, we use both height- and width-axial self-attention blocks in the U-net architecture, which calculate weights by incorporating the distances among protein atoms. Specifically, our model can break down into multiple modules: a U-net encoder which contains: *C*_128_-*A*_128_-*A*_128_-*A*_128_-*C*_256_-*A*_256_-*C*_512_-*C*_1024_-*C*_2048_-*C*_4096_, and a decoder consists of *CD*_2048_-*CD*_1024_-*CD*_512_-*CD*_256_-*CD*_128_. After the last layer in the decoder, a convolution is applied and followed by a Tanh to provide a predicted pairwise distance matrix 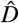 that captures the interaction between environmental protein atoms and the metal ion. The whole model is also elaborated in Figure 3 (b). The yellow, red, and blue blocks represent the features obtained after the convolutional layer, axial-attention layer, and skip connection, respectively. The labeled numbers indicate the dimensions of the hidden features.

#### Hyperparameters

The working environment is NVIDIA Quadro RTX 8000. In the training process, we choose Adam optimizer with β_1_ = 0.5 and β_2_ = 0.999. We trained our model for 60 epochs, the first 30 epochs have a learning rate of 0.0001, and it decays linearly in the following 30 epochs.

### Postprocessing

Given the coordinates of neighboring protein atoms and predicted pairwise distance matrix 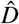, the location of the metal ion can be estimated. First, vectors located in the green region, in Figure 3 (c), are averaged *r* = (*r*_*row*_ + *r*_*column*_)*/*2, which represents the estimated distance between each atom in protein and the metal ion. The coordinates of metal ion and the *i*-th atom in protein are *x* = (*x*_1_, *x*_2_, *x*_3_)^*T*^ and *p*_*i*_ = (*p*_*i*,1_, *p*_*i*,2_, *p*_*i*,3_)^*T*^, respectively. Then the Euclidean distance between them is

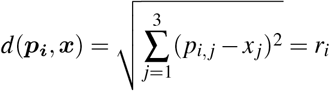

where *r*_*i*_ is the *i*-th element in *r*.

Thus, the difference of squared distance with respect to the baseline *p*_**1**_ becomes 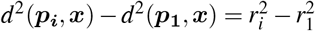, which can be reformulated as:

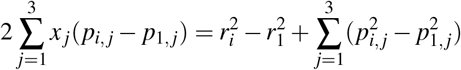

for *i*=2,3,…, 255.

In a matrix form, it can be rewritten as

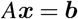

where *A* =[2(*p*_*i, j*_ − *p*_1, *j*_)]_*i*=2,3,…,255; *j*=1,2,3_ and the *i*-th element of *b* is 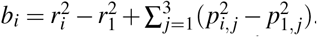. The best estimation of ligand coordinates can be obtained by minimizing mean squared error, in the following form:

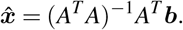

## Results

Table 1 displays the performance of various models, evaluated using two criteria: Euclidean distance between the predicted and actual locations, and the proportion of predicted locations that fall within a specified search region. Two commonly used thresholds are 2 Å and 4 Å. To ensure a fair comparison, BioMetAll and AutoDock Vina (37) use the same center of the local region that was used during data augmentation, investigating potential binding within a sphere of a 10 Å radius, which is the minimal search region in our model (Figure 2). Narrowing the search region can lead to improved results due to the integration of additional prior knowledge, thus a 10-angstrom search area is suggested to yield the most optimal results. Wang et al. choose a search radius of 18 Å after fine-tuning. Besides, BioMetAll yields multiple putative metal coordinates, and the group center includes the most candidate probes is chosen for each instance. Furthermore, grid sizes of 1 and 0.5 have been explored, as they do not exhibit a significant difference and the latter is considerably more time-intensive, the presented results utilize the default grid size 1. Our model was selected based on validation performance and tested on augmented testing data with 49,300 instances.

**Table 1.**
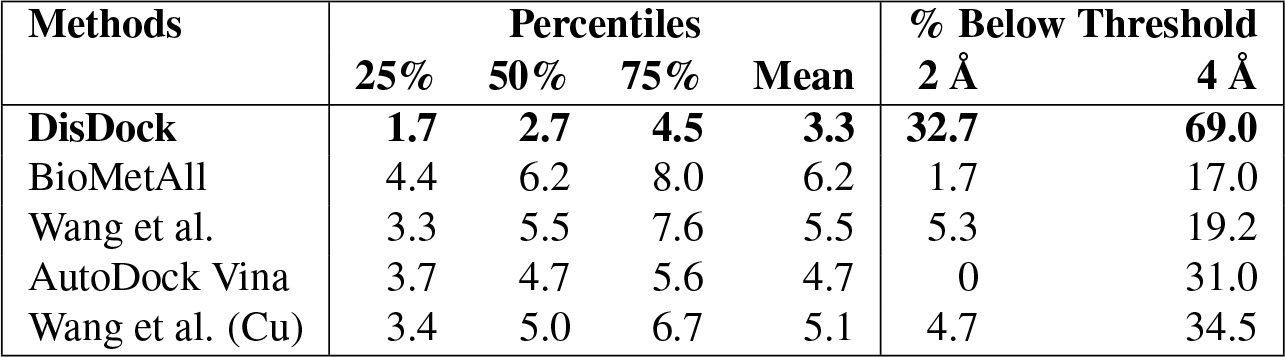
Performance comparison is conducted across various methods, with a focus on predictions that are within a Euclidean distance of less than 10 Å from the true location. AutoDock Vina: PDB files are handled through multiple preprocessing steps, and 34% of testing data are applicable and shown. More details of AutoDock Vina and related tools can be found in the document (33; 34; 35; 36). Wang et al. (Cu): While their model can be applied to all metal ions, it is solely trained on the copper ion. Therefore, model performances on the entire set and copper subset are shown on the third and last row, respectively.

| Methods | Percentiles |  |  |  | % Below Threshold |  |
| --- | --- | --- | --- | --- | --- | --- |
|  | 25% | 50% | 75% | Mean | 2 Å | 4 Å |
| **DisDock** | **1.7** | **2.7** | **4.5** | **3.3** | **32.7** | **69.0** |
| BioMetAll | 4.4 | 6.2 | 8.0 | 6.2 | 1.7 | 17.0 |
| Wang et al. | 3.3 | 5.5 | 7.6 | 5.5 | 5.3 | 19.2 |
| AutoDock Vina | 3.7 | 4.7 | 5.6 | 4.7 | 0 | 31.0 |
| Wang et al. (Cu) | 3.4 | 5.0 | 6.7 | 5.1 | 4.7 | 34.5 |

The results for the three methods evaluated on all augmented testing data are shown in the first three rows. Our DisDock model demonstrates better performance by achieving higher precision compared to the other models, and the median and mean Euclidean distance are 2.7 Å and 3.3 Å, respectively. Using commonly used thresholds of 2 Å and 4 Å as measures for success, our model successfully redock about 30% and 70% of cases, respectively, while other models have rates of less than 20%. Regarding AutoDock Vina, which is designed for docking with multi-atom ligands, not all metal-binding complexes can be properly preprocessed. In the fourth row, the results are limited to 34% of the testing data that remain after preprocessing. In addition, Wang et al. trained their model specifically for copper ions, so the last row illustrates their model performance on a subset of all copper ions. For these two subsets, our model maintains predictive power as it is in the entire data and still outperforms the other models. Detailed measurements are excluded here, with the full table accessible in Supporting information S3 Table.

Figure 5 demonstrates the performance of the DisDock model for predicting the precise location of *Zn*^2+^ on chain C of 1PL8. The actual location of the metal ion is indicated by the red point. Utilizing DisDock, three predictions were carried out, each with a distinct initial location and search region. These search regions are depicted as yellow (centered on residue ASER at position 46), pink (centered on residue LYS at position 344), and green (centered on residue GLU at position 155) for cases 1 through 3, while the corresponding initial metal locations are represented as gray dots in the figure. Despite the relatively large search regions encompassing neighboring atoms of the centered residues, the DisDock model achieves accurate predictions for the precise location of *Zn*^2+^ in all cases. Notably, the model’s performance remains consistent irrespective of the chosen initial locations and search regions.

**Fig 5.**
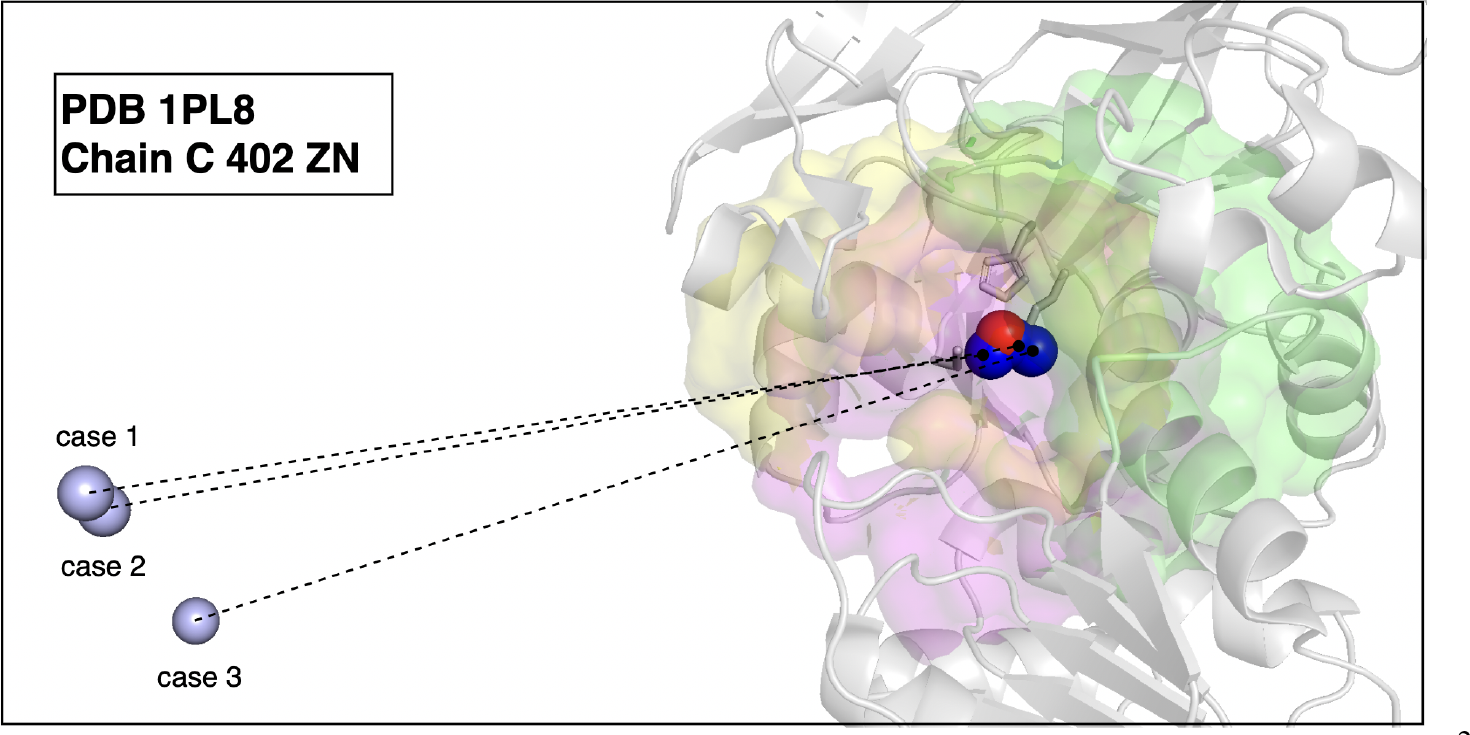
An example of redocking shown by 1PL8, specifically chain C at position 402 with *Zn*^2+^. The true location of the metal ion is depicted in red, while the blue and gray dots represent the predicted and initial locations, respectively. Three predictions, referred to as cases 1 through 3, were conducted using different initial locations and search environments. These search environments were centered around three residues: ASER at position 46 (highlighted in yellow), LYS at position 344 (highlighted in pink), and GLU at position 155 (highlighted in green). The colors in the shared area are blended together.

## Discussion

In this study, we developed DisDock, a deep learning method to predict protein-metal binding complex structures given the protein structure and the type of metal ion. DisDock has achieved significantly better performance than existing methods in terms of two commonly used metrics on a large dataset. DisDock combines the U-net architecture with self-attention modules, thereby more effectively encoding geometric features using atom-atom distance information as input. DisDock aims to learn the atom-atom distance matrix of the true protein-metal complex, mimicking the real protein-metal binding process. In this process, the metal ion starts from a location away from the true binding position and “finds” its way to the true binding position driven by the physical interactions between the metal ion and protein atoms around the binding site. By mimicking the physical process, our model therefore may learn the parameters that resemble the actual physical interactions among the atoms. Future works that analyze the learned parameters may shed lights on the real physics of protein-metal binding.

Metal1D and Metal3D mentioned earlier take the entire protein as input and only predict zinc binding. Therefore, we did not compare our method with Metal1D and Metal3D. Moreover, certain techniques developed for multi-atom ligands (37; 38) also fall short when applied to metal ions. For instance, GNINA (38) excludes ligands with weights less than 150 Da during training, and most metal ions are lighter than 150 Da. Considering docking engines, such as AutoDock Vina, that require multiple preprocessing steps, the proper handling of all metal-binding complexes becomes a challenge. It is even more complicated when taking experimental structures (for instance, from the Protein Data Bank) as input because waters, co-factors, and ions seemed unnecessary for the docking need to be removed beforehand. Then receptors and ions need to be prepared separately.

In this study, we have generated the data with the knowledge of the protein-metal binding complex structures. While being a limitation for the current setting, the framework has the potential to handle the flexibility of proteins, thanks to the computational efficiency gained by including only a subset of protein atoms. By augmenting the data to generate local regions partially overlapping with the true local region formed by the subset of atoms closest to the metal ion, the augmented data resembles the realistic scenario where we predict the local region and start from a predicted local region. Our result showed that even if starting from a local region partially overlaps with the true local region, very satisfactory performance can be achieved, indicating that an end-to-end pipeline can be successfully built. This pipeline will start from whole proteins by first predicting a subset of atoms close to the metal binding site and then using that subset of atoms to predict the protein-metal binding. Some methods have been developed for this purpose, which can be used to build the end-to-end pipeline.

Future work will focus on investigating increasingly complex scenarios to advance the accuracy and capacity for generalization of the technique. This includes exploring the docking of multi-atom ligands, accommodating the flexibility of proteins during the docking process, and tackling blind docking challenges. Furthermore, the exploration of cutting-edge architectures will be conducted to enhance the capabilities of molecular docking algorithms and elevate their performance in a wide range of molecular docking scenarios.

## Supporting information

**S1 Table.**
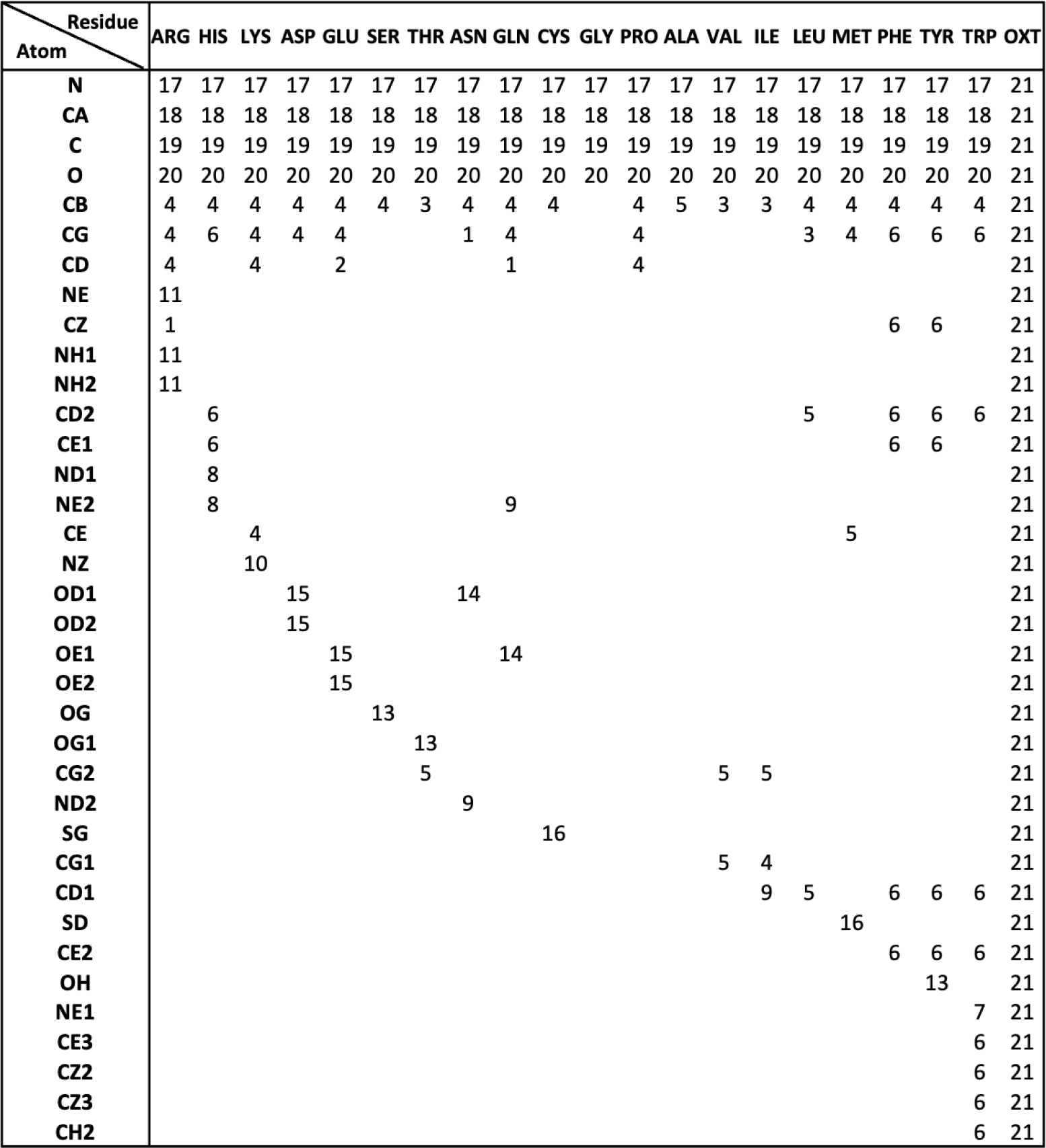
Specification of protein atom channel. When generating input of size 23 × 256 × 256, each environment atom is encoded based on its amino acid residue and type. For instance, the rows and columns corresponding to nitrogen will be assigned a value of 1 on the 17th channel if this nitrogen atom comes from residues ARG.

**S2 Table.** Metal ions investigated in this study and their sample sizes in the training, validation, and test data.

| Metal ions | train | val | test | total |
| --- | --- | --- | --- | --- |
| MG | 8273 | 1418 | 2723 | 12414 |
| ZN | 8550 | 1061 | 2236 | 11847 |
| CA | 6570 | 649 | 2122 | 9341 |
| NA | 3267 | 382 | 1296 | 4945 |
| MN | 2555 | 732 | 465 | 3752 |
| K | 1980 | 200 | 443 | 2623 |
| FE | 721 | 122 | 175 | 1018 |
| NI | 537 | 148 | 55 | 740 |
| CO | 399 | 53 | 124 | 576 |
| FE2 | 341 | 39 | 167 | 547 |
| CU | 207 | 46 | 25 | 278 |
| BA | 66 | 1 | 7 | 74 |
| MO | 18 | 16 | 26 | 60 |
| W | 36 | 16 | 0 | 52 |
| CU1 | 8 | 9 | 2 | 19 |
| CD | 5 | 1 | 0 | 6 |
| Total | 33533 | 4893 | 9866 | 48292 |

**S1 Fig.**
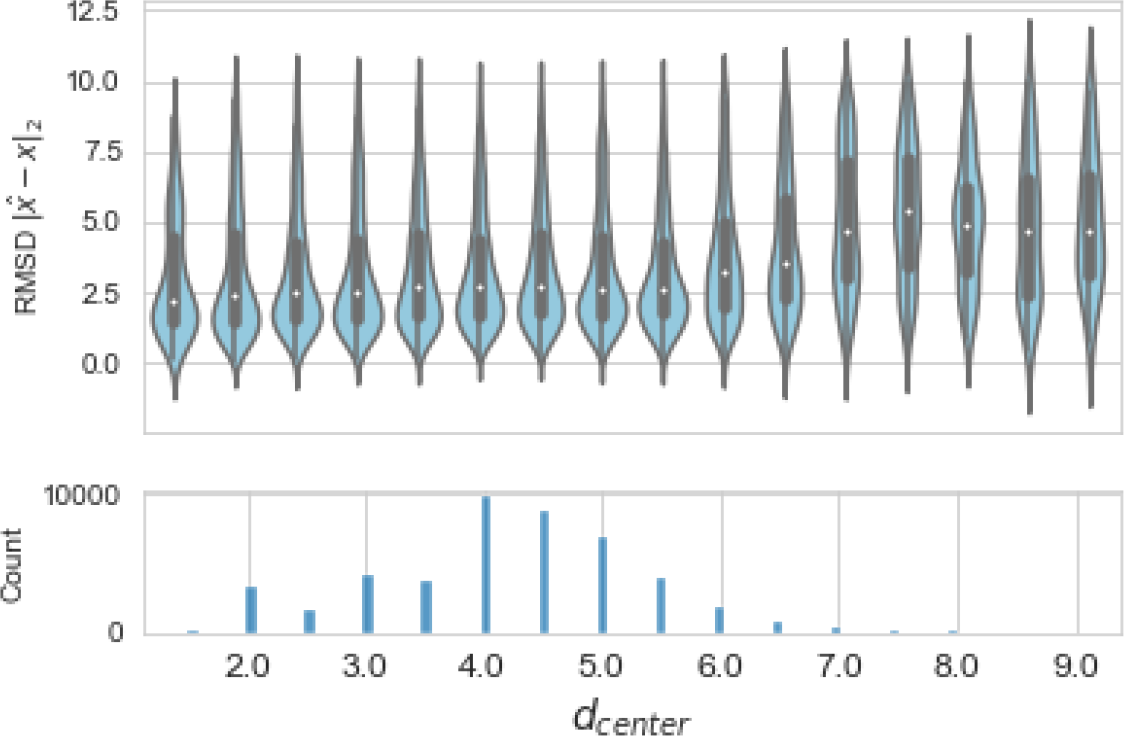
Model performance by distance from metal ion. Figure illustrates the performance of our model with respect to translations of the environment center relative to the metal ion. The x-axis of the plot shows the Euclidean distance between the metal ion and the augmentation center, represented by *d*_*center*_ and divided into discrete intervals. The y-axis of the top panel represents the precision of the model’s predictions, and the bottom panel shows a histogram of the number of instances. Our model performs well when *d*_*center*_ is less than or equal to 6 Å, with a mean and median around 2.5 Å or less. However, as the augmentation center moves away from the metal ion, the task of accurately predicting the true metal binding conformation becomes more challenging..

**S2 Fig.**
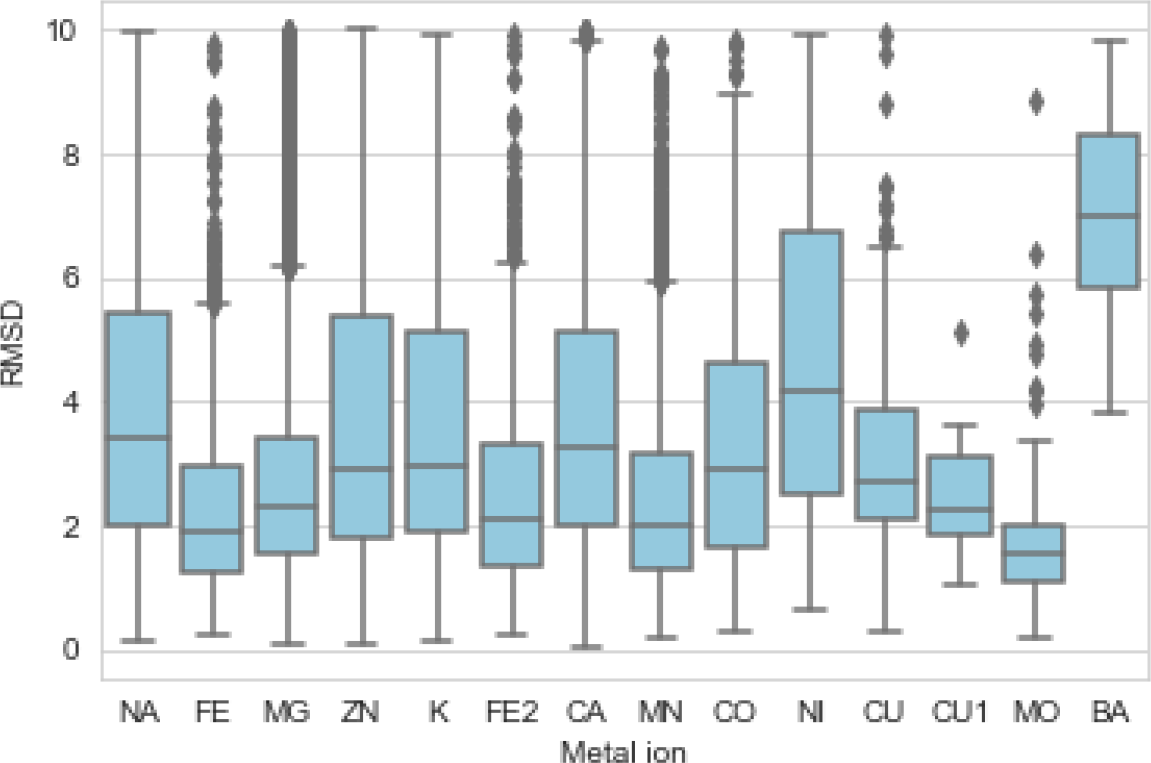
Model performance by metal ion type. The performance of the model for different types of metal ions is demonstrated in Figure. The horizontal axis of the figure shows 14 types of metal ions, excluding *W* ^6+^ and *Cd*^2+^ which were not present in the testing data. The vertical axis shows the conformational precision measured in Euclidean distance. While the prediction accuracy varies across different types of metal ions, the majority of them have a median Euclidean distance of less than 3 Å.

**S3 Table.** A detailed performance comparison across various computational methods. The DisDock method exhibits notably superior performance in prediction precision across multiple metrics, including different percentile measures and percentages below given distance thresholds, relative to other methods. Two additional subset performances are included: results utilizing DisDock on AutoDock Vina processed testing data, and results specifically for copper ions.

| Methods | Percentiles |  |  |  | % Below Threshold |  |
| --- | --- | --- | --- | --- | --- | --- |
|  | 25% | 50% | 75% | Mean | 2 Å | 4 Å |
| **DisDock** | **1.7** | **2.7** | **4.5** | **3.3** | **32.7** | **69.0** |
| BioMetAll | 4.4 | 6.2 | 8.0 | 6.2 | 1.7 | 17.0 |
| Wang et al. | 3.3 | 5.5 | 7.6 | 5.5 | 5.3 | 19.2 |
| **DisDock** | **1.6** | **2.4** | **3.9** | **3.0** | **37.6** | **75.6** |
| AutoDock Vina | 3.7 | 4.7 | 5.6 | 4.7 | 0 | 31.0 |
| **DisDock** | **2.1** | **2.6** | **3.8** | **3.3** | **20.1** | **76.1** |
| Wang et al. (Cu) | 3.4 | 5.0 | 6.7 | 5.1 | 4.7 | 34.5 |

